# Automated Gleason grading of prostate cancer tissue microarrays via deep learning

**DOI:** 10.1101/280024

**Authors:** Eirini Arvaniti, Kim S. Fricker, Michael Moret, Niels J. Rupp, Thomas Hermanns, Christian Fankhauser, Norbert Wey, Peter J. Wild, Jan H. Rueschoff, Manfred Claassen

## Abstract

The Gleason grading system remains the most powerful prognostic predictor for patients with prostate cancer since the 1960’s. Its application requires highly-trained pathologists, is tedious and yet suffers from limited inter-pathologist reproducibility, especially for the intermediate Gleason score 7. Automated annotation procedures constitute a viable solution to remedy these limitations.

In this study, we present a deep learning approach for automated Gleason grading of prostate cancer tissue microarrays with Hematoxylin and Eosin (H&E) staining. Our system was trained using detailed Gleason annotations on a discovery cohort of 641 patients and was then evaluated on an independent test cohort of 245 patients annotated by two pathologists. On the test cohort, the inter-annotator agreements between the model and each pathologist, quantified via Cohen’s quadratic kappa statistic, were 0.75 and 0.71 respectively, comparable with the inter-pathologist agreement (kappa=0.71). Furthermore, the model’s Gleason score assignments achieved pathology expert-level stratification of patients into prognostically distinct groups, on the basis of disease-specific survival data available for the test cohort.

Overall, our study shows promising results regarding the applicability of deep learning-based solutions towards more objective and reproducible prostate cancer grading, especially for cases with heterogeneous Gleason patterns.

## Introduction

Prostate cancer is the second leading cause of cancer death in men^1^. Prostatic carcinomas are graded according to the Gleason scoring system which was first established by Donald Gleason in 1966^2^. The Gleason scoring system is acknowledged by the World Health Organization (WHO) and has been modified and revised in 2005 and 2014 by the International Society of Urological Pathology (ISUP)^3^. Despite several changes in clinical diagnosis of prostate cancer, the histological Gleason scoring system is still the most powerful prognostic tool^4^. The assessment is based exclusively on the architectural pattern of the tumour, i.e. the Gleason patterns. Different histological patterns are assigned numbers from 1 (well differentiated) to 5 (poorly differentiated). Gleason pattern 3 describes well-formed, separated glands, variable in size. Gleason pattern 4 includes fused glands, cribriform and glomeruloid structures and poorly formed glands. Gleason pattern 5 involves poorly differentiated individual cells, sheets of tumour, solid nests, cords and linear arrays as well as comedonecrosis. The final Gleason score is reported as the sum of the two most predominant patterns present in the histological specimen, and in current clinical practice the lowest Gleason score assigned is Gleason 6 (3+3)^5^.

Immunohistochemistry is routinely generated in clinical laboratories with good reproducibility. However, the final Gleason score annotation of the stained tissue slides is dependent on the evaluation of the respective pathologist, who thus constitutes an important factor for stratification and therapeutic decisions. The histological assessment of human tissue—based on visual, microscopy-based evaluation of non-trivial cellular and morphological patterns—is time-consuming and often suffers from limited reproducibility^6^. For prostate cancer in particular, intermediate-risk Gleason patterns 3 and 4 can be very difficult to assign unambiguously.

Automated computational approaches operating on digital pathology images bear the potential to overcome the above-mentioned limitations, deliver reproducible results and achieve high throughput by multiplexing of computational resources^7^. Earlier computational approaches developed for this purpose build on explicit, *a priori* defined image features and employ conventional regression or classification techniques to perform feature selection and association with clinical parameters^8–10^.

In recent years, deep learning has emerged as a disruptive alternative to the aforementioned feature engineering-based techniques. Deep learning systems rely on multi-layered neural networks that are able to extract increasingly complex, task-related features directly from the data. Recent developments in neural network architecture design and training have enabled researchers to solve previously intractable learning tasks in the field of computer vision^11^. As a result, deep learning-based approaches have become very successful in addressing a wide range of biomedical image analysis tasks^12-15^.

Recent studies (e.g. ^16–19^) have shown that deep learning systems can accurately detect malignancies in histopathological images. Prior work on the analysis of prostate cancer digital pathology images includes detection of cancerous tissue^18^, prediction of *SPOP* mutation status^20^ and of cancer recurrence^21^, as well as tissue heterogeneity characterization via unsupervised learning^22^. A first deep learning-based approach to Gleason score prediction is the study of Källén *et al.*^23^. However, the assessment of their method was limited to tissue slides with homogeneous Gleason grading, despite the fact that, typically, tissue slides contain heterogeneous Gleason pattern regions. In a more recent work, Zhou *et al.* focused on intermediate Gleason scores^24^. Their algorithm was tested on whole slide images from The Cancer Genome Atlas (TCGA)^25^ achieving an overall accuracy of 75% in differentiating Gleason 3+4 from Gleason 4+3 slides. Finally, del Toro *et al.* also used prostate cancer whole slide images from TCGA and trained a binary classifier to discriminate low (7 or lower) versus high (8 or higher) Gleason score images^26^. The above studies suggest that automated Gleason grading via deep learning is a feasible task, but were all evaluated in a limited setting—either on images with homogeneous Gleason patterns or as a binary classification problem on a limited set of Gleason scores.

## Results

In this study, we focus on a well annotated dataset of prostate cancer tissue microarrays^27^ and demonstrate that a convolutional neural network can be successfully trained as a Gleason score annotator. As opposed to previous studies, we both train and evaluate our model on the basis of detailed manual expert Gleason annotations of *image subregions* within each TMA spot image. Given the relatively small size of our training dataset (641 patients), we find transfer learning, strong regularization and balanced mini-batches to be crucial for successfully training the classifier. We then evaluate the model’s predictive performance on a separate test cohort of 245 patients, independently annotated by two specifically trained uropathologists (K.S.F., J.H.R.). This design allows us to quantify the inter-pathologist variability on this particular test cohort and compare it with the neural network performance. Additionally, we have benchmarked the deep learning approach against the two pathologists on a survival analysis task. This task allowed an objective comparison since survival data constitutes a precisely measurable ground truth as opposed to Gleason annotations which can be subjective. We show that the Gleason score groups assigned by the neural network achieve pathologist-level risk stratification of the prostate cancer patients in our test cohort.

### Tissue microarray resource with Gleason score annotated subregions

Our dataset comprises five tissue microarrays (TMAs), each containing 200-300 spots. Spots containing artefacts or non-prostate tissue (e.g. lymph node metastasis) were excluded from the study. The prostate TMA spots were annotated by a first pathologist (K.S.F.) by carefully delineating cancerous regions and assigning a Gleason pattern of 3, 4 or 5 to each region. Examples of annotated regions are depicted in **Figure 1a**. TMA spots without any cancerous region were marked as benign. The distribution of Gleason scores across different tissue microarrays is summarized in **Table 1**. TMA 80 contains the highest number of cases and was chosen as the test cohort. TMA 76 was chosen as the validation cohort because it contains the most balanced distribution of Gleason scores. The other three TMAs were used as training cohort. The TMA spots in the test cohort were independently annotated by a second pathologist (J.H.R.) to allow for quantification of inter-pathologist variability.

**Table 1:**
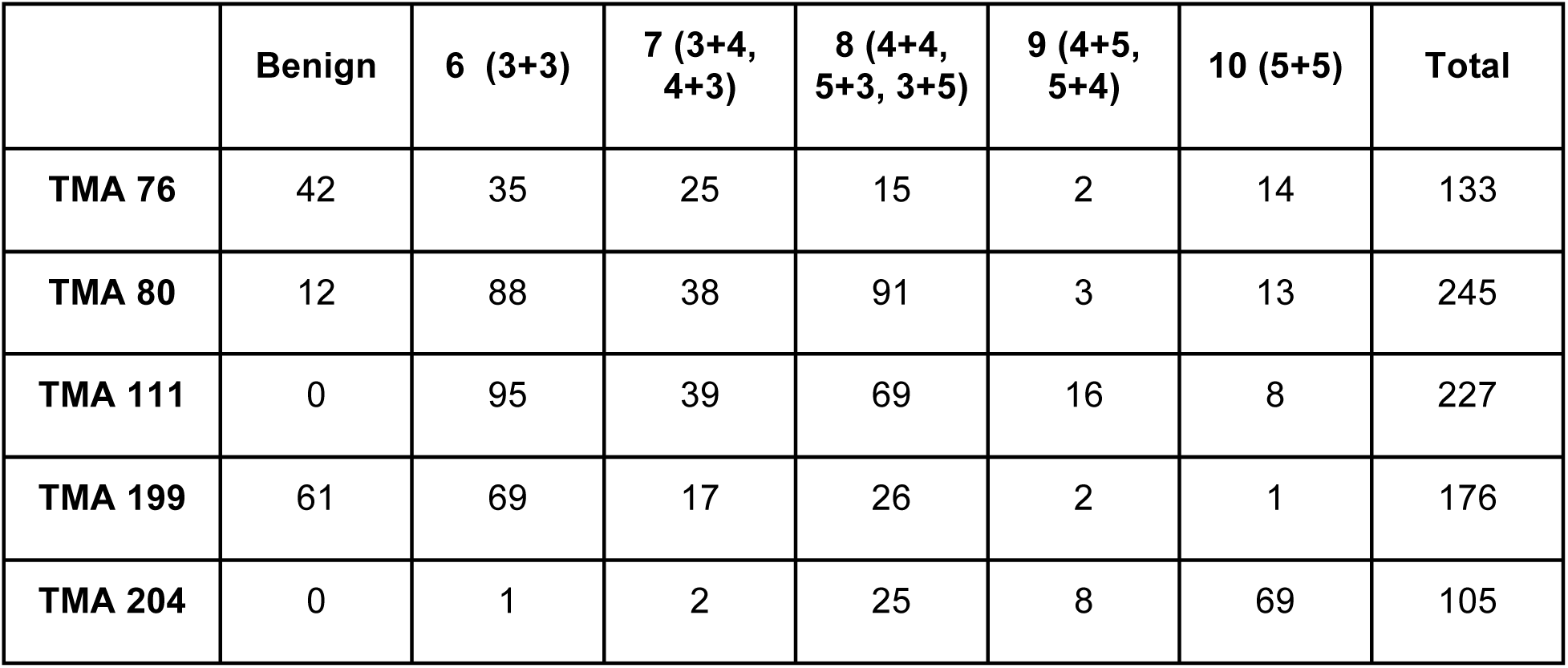
Dataset Gleason annotation summary. Tissue Microarrays 111, 199, 204 were used as training set, TMA 76 as validation set and TMA 80 as test set.

**Figure 1:**
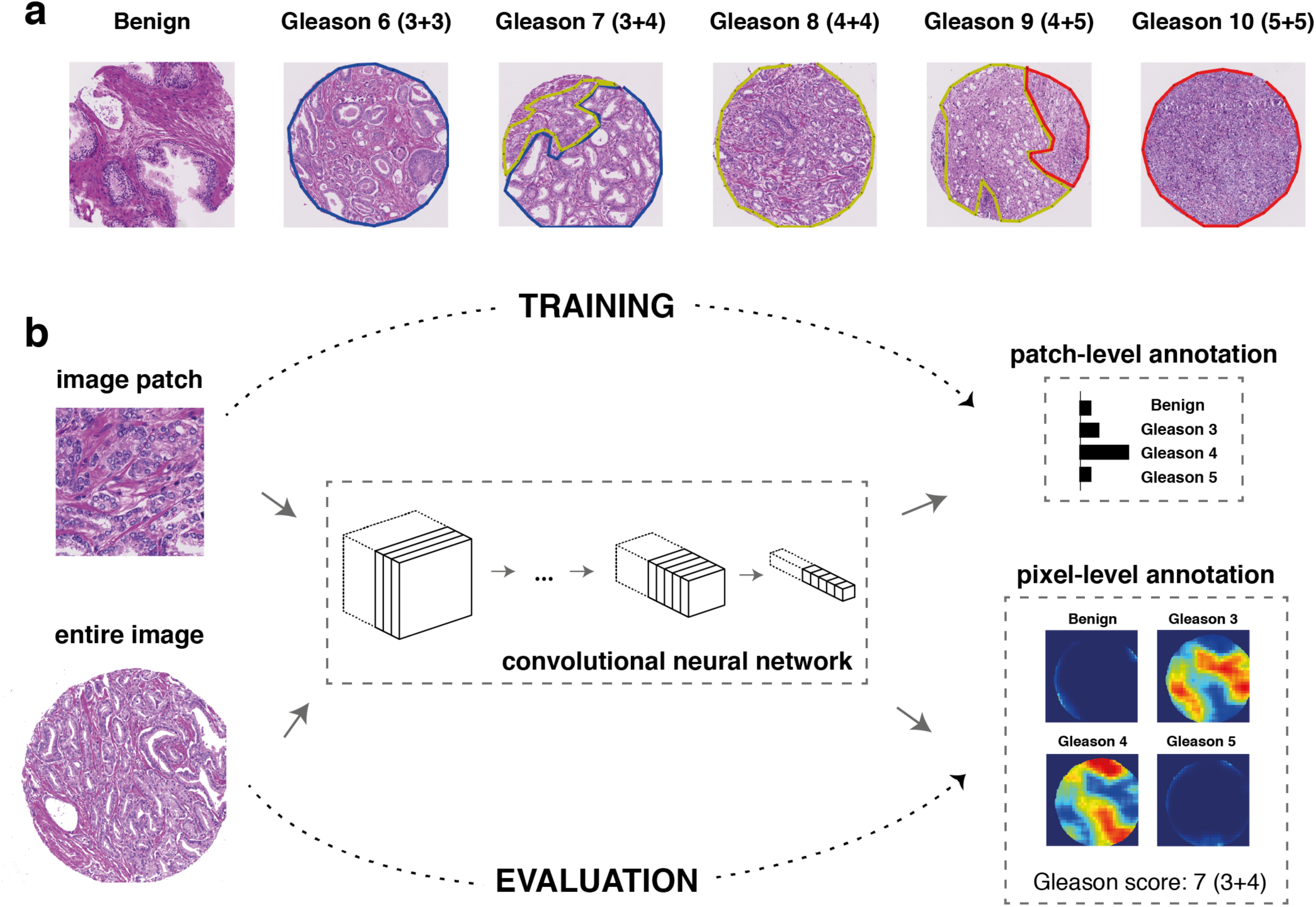
Overall annotation procedure. **(a)** Examples of TMA spot Gleason annotations provided by the pathologists (blue: Gleason pattern 3 region, yellow: Gleason pattern 4 region, red: Gleason pattern 5 region). **(b)** During the training phase (top row), a deep neural network was trained as a patch-level classifier. We used the MobileNet architecture, whose main building blocks are “depthwise separable” convolutions: a special type of convolution block with considerably fewer parameters than normal convolutions. Convolution blocks are used to extract increasingly complex features from the input image. Following the convolution blocks, a global average pooling layer computes the spatial average of each feature map at the last convolution layer, effectively summarizing the locally-detected patterns across the entire image. Finally, the output layer produced the final classification decision for each input image patch by computing a probability distribution over the four Gleason classes considered in this study. During the evaluation phase (bottom row), the trained patch-level convolutional neural network was applied to entire TMA spot images in a sliding window fashion and generated pixel-level probability maps for each class. A Gleason score was assigned to a TMA spot as the sum of the primary and secondary Gleason patterns detected (above a threshold) in the corresponding output pixel-level maps.

In addition to Gleason score annotations, clinical data including survival information was available for three of the TMAs^27^. Prostate cancer is known to have a favorable prognosis compared to other types of cancer, and therefore relatively few death events are present in the dataset. Restricted to TMA spots considered in this study, TMA 80 has the highest number of death events (n = 30), followed by TMA 111 (n = 7) and TMA 76 (n = 0).

### Automated Gleason score annotation via deep learning

Small image regions (image “patches”) were extracted from benign tissue and the cancer annotated regions and used to train a patch-based classifier (**Fig. 1b**). Once trained, the patch-level classifier can be easily converted to a pixel-level annotator (details in **Methods**) and, therefore, can be used to assign Gleason scores to entire TMA spot images (**Fig. 1b**). To choose a suitable classifier, we explored different convolutional neural network architectures which have shown excellent performance on the ImageNet competition^28^, namely VGG-16^29^, Inception-V3^30^, ResNet-50^31^, DenseNet-121^32^ and MobileNet^33^. As expected, given the limited size of our dataset, fine-tuning the networks pretrained on ImageNet achieved better performance on the validation set than training from scratch **(Supplementary Fig. S1)**. Details about the patch generation procedure and model training can be found in **Methods**.

In our benchmark, the best performing network architecture was MobileNet (using width multiplier *α* = 0.5). MobileNets are designed as small models able to be trained on mobile devices and, in this application, where dataset size is limited and feature space is restricted in comparison to natural images, it turned out that the relatively low number of parameters helped to avoid severe overfitting without causing a drop in performance. In the validation cohort, MobileNet achieved a macro-average recall of 70% for classifying patches as benign, Gleason 3, Gleason 4 or Gleason 5. Specifically for each class, the recall values were: benign 63%, Gleason 3 72%, Gleason 4 58%, Gleason 5 88%. The full confusion matrix is shown in **Supplementary Figure S2**. In this cohort, the value of Cohen’s quadratic kappa^34^ for patch-level classification was 0.67.

We then evaluated the model’s predictions at the patch level in the test cohort (**Fig. 2**). The results of comparing the model’s annotations with annotations from the first pathologist (who has also annotated the training cohort) showed that, although there is no perfect agreement, most mis-classifications were found within neighbouring Gleason patterns (**Fig. 2a**). For instance, 31% of patches annotated as Gleason 5 were predicted as Gleason 4, but only 5% were predicted as lower Gleason patterns. For this comparison, Cohen’s quadratic kappa was 0.55 and macro-average recall was 0.58. Comparing the model’s annotations with annotations from the second pathologist, we observe a similar agreement pattern, resulting in a kappa value of 0.49 and macro-average recall of 0.53 (**Fig. 2b)**. Finally, we quantified the inter-pathologist agreement on the test cohort (kappa = 0.67, macro-average recall = 0.71 considering the first pathologist’s annotations as ground truth), which is at higher but still comparable levels as the agreement between model and pathologist annotations (**Fig. 2c**). The degree of overlap between model and pathologist annotations was also visually depicted via Venn diagrams in **Figure 2d**.

**Figure 2:**
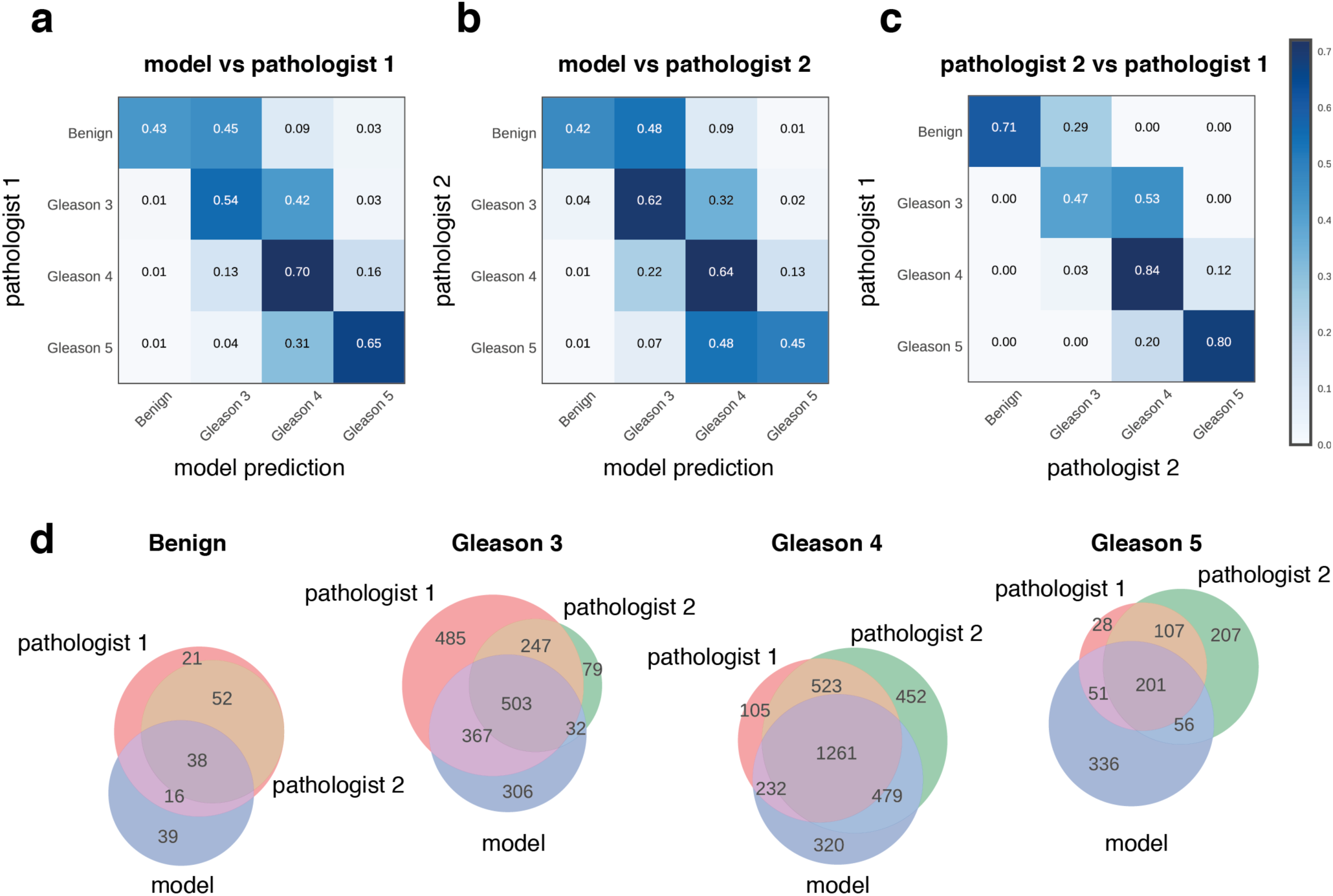
Model evaluation on test cohort (image patch level) and inter-pathologist variability. All confusion matrices were normalized per row (ground truth label) reflecting the recall metric for each class. **(a)** Patch-based model annotations compared with annotations by 1st pathologist. **(b)** Patch-based model annotations compared with annotations by 2nd pathologist. **(c)** Annotations by 2nd pathologist compared with annotations by 1st pathologist. **(d)** Venn diagrams illustrating the overlap in patch-level Gleason annotations produced by the deep learning model and the two pathologists.

As a next step, we evaluated the model’s predictions at the TMA spot level. To this end, we generated pixel-level probability maps for each class by applying the trained neural network in a sliding window fashion (**Fig. 3**, for details see **Methods**). These pixel-level maps enable visual comparison with human annotations and can be easily evaluated by pathology experts. For instance, the TMA spot in **Figure 3a** was a clear Gleason 6 (3+3) case according to both pathologists and model annotations. We also observed that the model annotated the upper region of this TMA spot as benign, in accordance with the pathologists, except for a small part that was annotated as Gleason pattern 3. Interestingly, retrospective assessment of this part by the pathologists confirmed the presence of a small focus of atypical glands. The TMA spot in **Figure 3b** was annotated as Gleason 8 (4+4) by the two pathologists and the model. Here we observed some annotation errors made by the model in regions close to tissue borders, often affected by tissue preparation artefacts. Such small mis-labeled regions do not influence the final Gleason score assignment for entire TMAs, as we only take into account detected patterns that exceed a predefined threshold (details in **Methods**). As a further example, the TMA spot presented in **Figure 3c** was annotated as Gleason 8 (4+4) by the first pathologist and as Gleason 6 (3+3) by the second. The model annotations split the TMA spot image into distinct Gleason pattern 3 and Gleason pattern 4 regions, resulting in a Gleason score 7 (4+3) final assignment. Similarly for the TMA spot presented in **Figure 3d**, which was annotated as Gleason 8 (4+4) by the first pathologist and as Gleason 10 (5+5) by the second. The model annotations split the TMA spot image into distinct Gleason pattern 4 and Gleason pattern 5 regions, resulting in a Gleason score 9 (5+4) final assignment.

**Figure 3:**
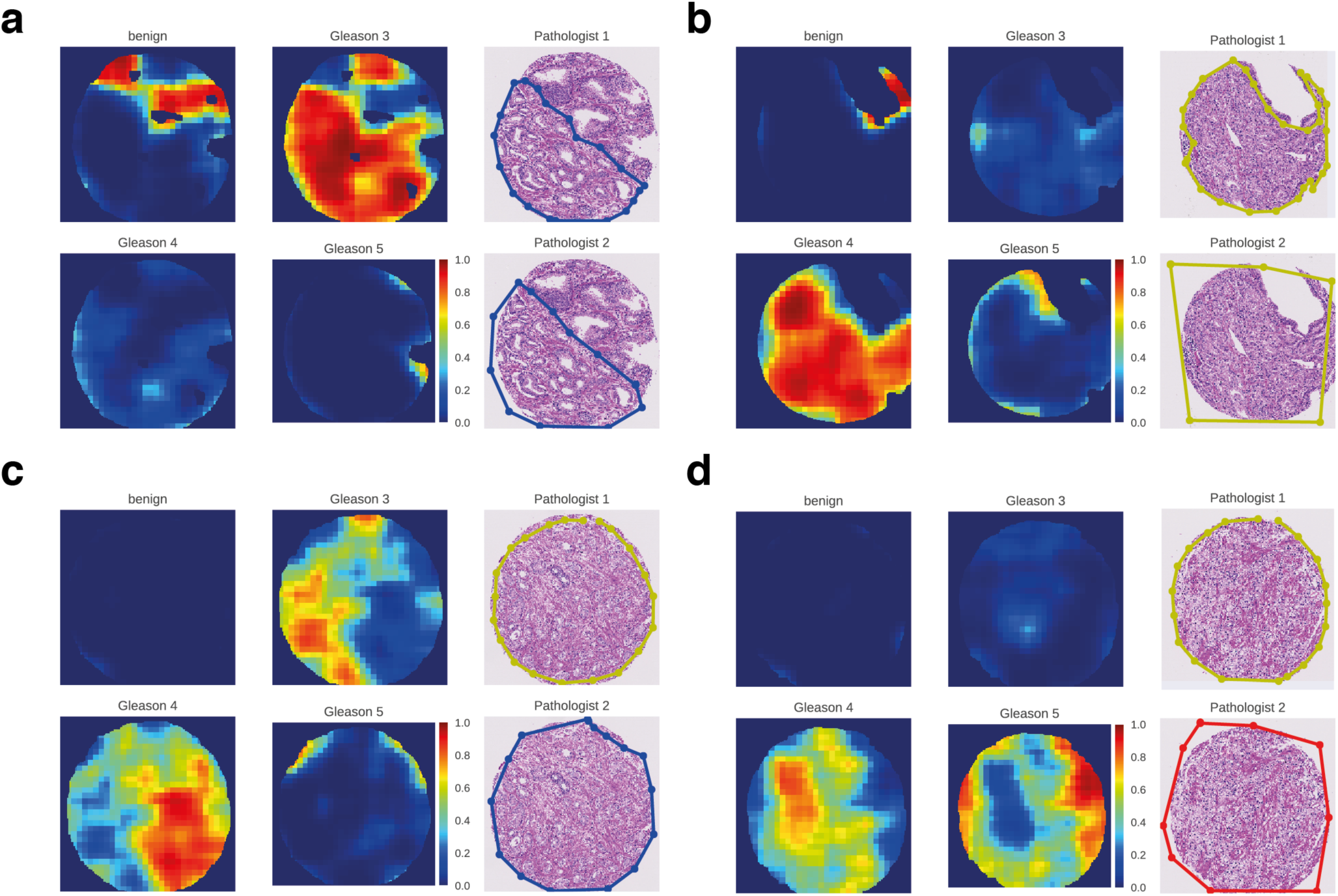
Representative examples of model predictions as pixel-level probability maps and visual comparison with pathologist annotations. Each subfigure **(a-d)** corresponds to a different TMA spot. Within each subfigure **(a-d),** the subplots in the right-most column show the Gleason patterns assigned by the two pathologists (blue: Gleason 3 region, yellow: Gleason 4 region, red: Gleason 5 region). The other four subplots show the model’s Gleason annotations.

Summarizing the pixel-level Gleason annotation maps produced by either the pathologists or the model (detailed description in **Methods**), a Gleason score in the range 6 - 10 was assigned to each TMA spot, as the sum of the two most predominant Gleason patterns. If no cancer was detected, the TMA spot was classified as benign. For this task, the inter-annotator agreement between the model and each pathologist reached kappa = 0.75 for the first pathologist and kappa = 0.71 for the second (**Fig. 4**). These values are at the same level as inter-pathologist agreement (kappa = 0.71).

**Figure 4:**
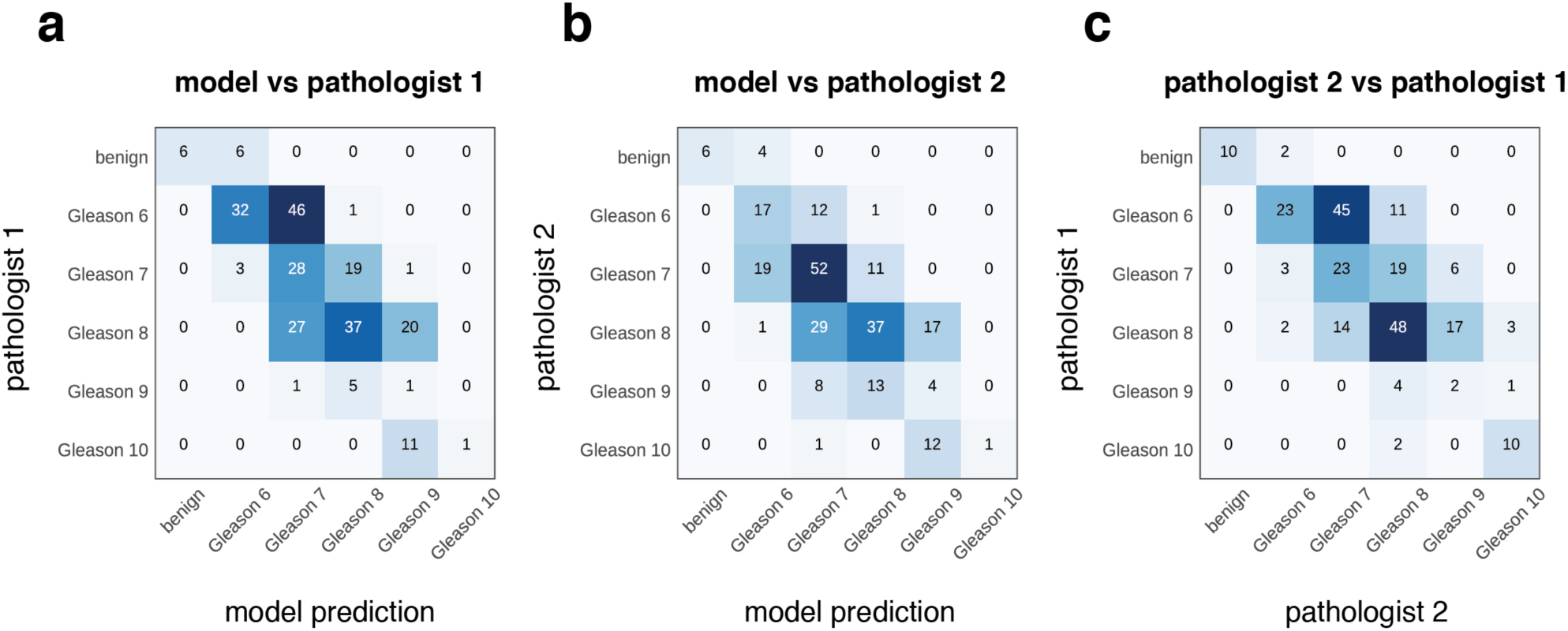
Model evaluation on test cohort (TMA spot level) and inter-pathologist variability. Each TMA spot is annotated with detected Gleason patterns (Gleason 3, 4 or 5) by the model and two pathologists. Then, a final Gleason score is assigned as the sum of the two most predominant Gleason patterns. If no cancer is detected, the TMA spot is classified as benign. We show confusion matrices for the comparison of Gleason score assignments by **(a)** the model and the first pathologist, **(b)** the model and the second pathologist, **(c)** the two pathologists.

### Model interpretation identifies specific morphological patterns as determinants of automated annotation performance

Complex machine learning models, such as deep neural networks, are often criticized by clinicians as non-interpretable black boxes. To provide insight and gain trust in the model predictions, we visually studied image patches that the model predicts correctly and with high confidence (**Fig. 5**). Furthermore, we evaluated morphological patterns utilized by the model for assigning Gleason scores and compared these patterns to the ones expected by a pathology expert. We observed clear differences in the gland architecture among patches annotated with different Gleason grades by the model. Prostate glands in benign patches contained a well-formed outer layer of basal cells and showed no evidence of cytological atypia. In Gleason 3 patches, glands were variable in size but still round-shaped and well-formed. In Gleason 4 patches, we observed merged glands, small in size and irregularly shaped, as well as implied cribriform pattern. Finally, in Gleason 5 patches we found mostly the absence of gland formation and solid sheets of tumour.

**Figure 5:**
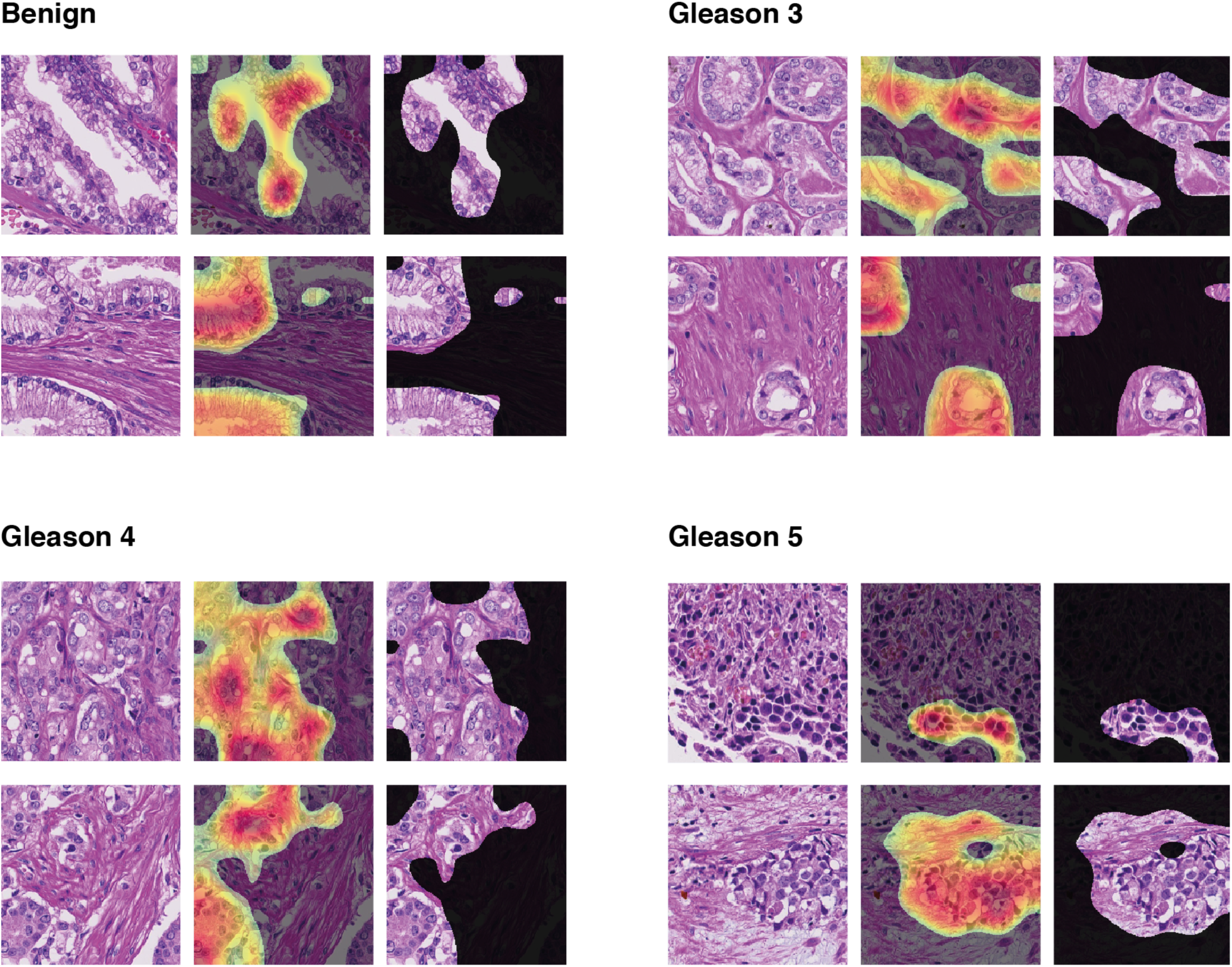
Model interpretation via class activation mapping^35^. For each class, we show two examples of image patches that were confidently and correctly classified by the deep learning model. In addition, the regions where the model is focusing on in order to make predictions are highlighted. In each example, the first column shows the image patch. In the second column, a heatmap generated by the class activation mapping technique is overlaid, highlighting the most important regions for the model predictions. In the third column, only the highlighted part of the image is shown.

To evaluate morphological patterns utilized by the annotation model, we employed the technique of class activation mapping (CAM)^35^ and highlighted class-specific discriminative parts of the confidently-predicted image patches. According to CAM analysis, the model is indeed focusing on the epithelium and ignoring stroma regions (**Fig. 5**). Especially for predicting the Gleason 3 pattern, we observed that the model focuses on gland junctions making sure that the glands are not fused, which would lead to a Gleason 4 pattern annotation.

### Automated Gleason score annotation yields pathology expert-level survival stratification

As a final step, for the test cohort, we studied survival stratification on the basis of the model annotations and those obtained by pathology experts. Patients were split into three risk groups on the basis of corresponding Gleason score assignments: low risk (Gleason ≤ 6), intermediate risk (Gleason 7), high risk (Gleason ≥ 8). Kaplan-Meier estimators of disease-specific survival showed differences for the individual risk groups considered (**Fig. 6a)**. Results about overall survival and biochemical recurrence-free survival followed similar trends (**Supplementary Fig. S3)**. We observed that the stratification achieved by the model for separating the low-risk and intermediate-risk groups was more significant (Benjamini Hochberg-corrected^36^ two-sample logrank p-value = 0.098) than the one achieved by either pathologist (BH-corrected two-sample logrank p-values = 0.79 and 0.292). The model automatically annotated Gleason 3 and 4 pattern regions and, eventually, assigned the heterogeneous Gleason score 7 in more cases than either pathologist. It is worth noting the inter-pathologist variability regarding Gleason score 7: for this intermediate-risk group, the two pathologists agreed on 19 cases and disagreed on 50 cases (**Fig. 6b**). In addition, we observed that 59% of the cases annotated as Gleason 7 by the first pathologist and, coincidentally, 59% of the cases annotated as Gleason 7 by the second pathologist were also annotated as Gleason 7 by the model. For the cases assigned to the low-risk group (Gleason ≤ 6) by the model, no disease-specific death event occurred. In contrast, for the cases labeled as low-risk by the first pathologist and second pathologist, 3 and 2 disease-specific death events occurred, respectively. These results demonstrate that the automated procedure achieved a pathologist-level survival stratification in this cohort of patients (see also **Supplementary Table S1)**.

**Figure 6:**
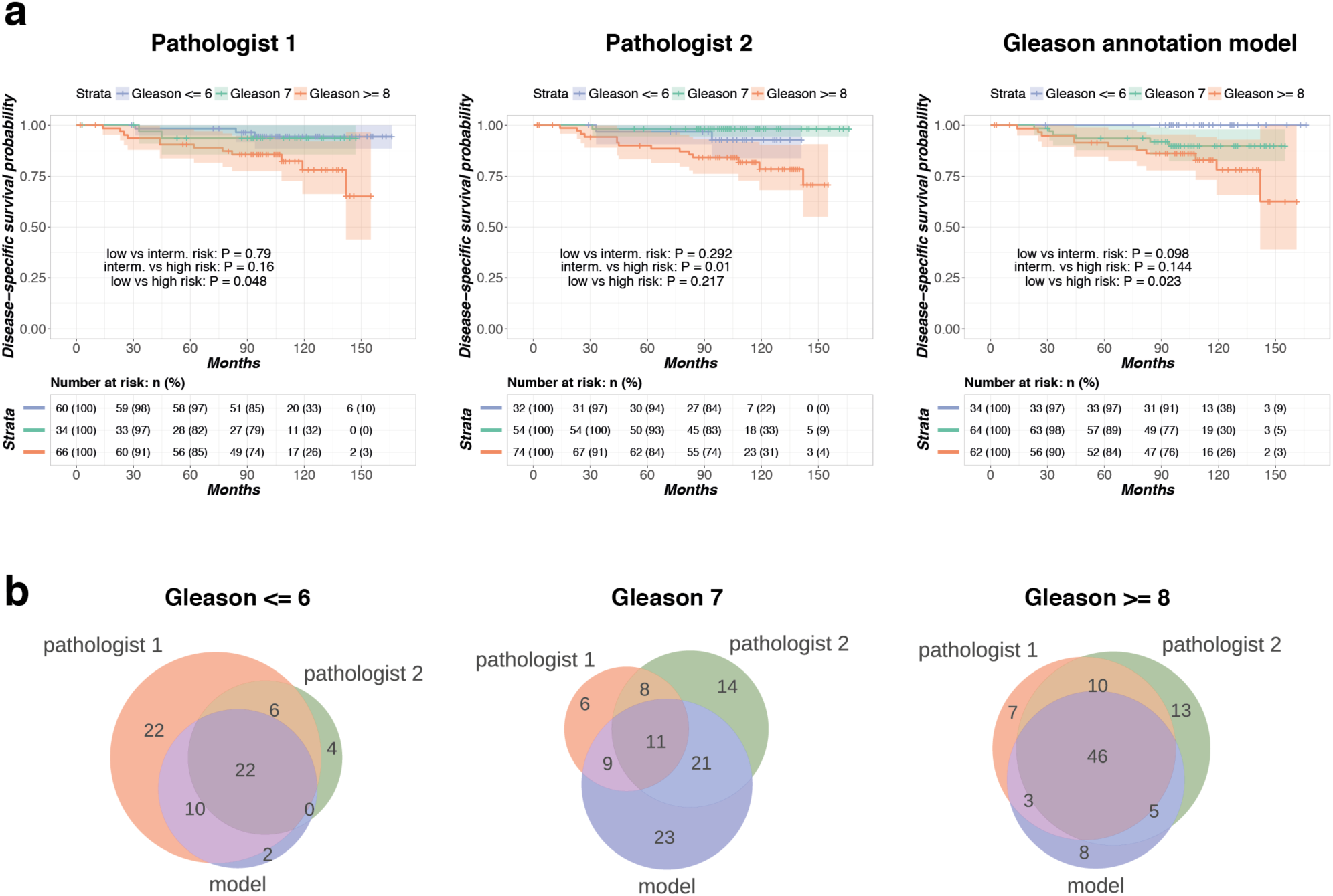
Disease-specific survival analysis results. **(a)** Kaplan-Meier curves for patients who were split into three risk groups according to Gleason score annotations by the model and two pathologists. The shaded regions indicate 95% confidence bands. P-values for pairwise two-tailed logrank tests with Benjamini-Hochberg correction are reported. **(b)** Venn diagrams illustrating overlap in model-based and pathologist annotation-based assignment of patients into Gleason score groups.

## Discussion

In this work, we have trained a convolutional neural network as Gleason score annotator and used the model’s predictions to assign patients into low, intermediate and high-risk groups, achieving pathology expert-level stratification results. The low-risk and intermediate-risk groups defined by the model’s predictions were more significantly separated compared to the corresponding groups defined by either pathologist’s annotations. To our knowledge, this is the first study where deep learning-based predictions are used for survival analysis in a prostate cancer cohort. Furthermore, visual inspection of morphological patterns obtained via the class activation mapping technique confirmed that the network is capturing class-discriminative information on prostate gland architecture for assigning Gleason scores.

This work is affected by some limitations, motivating future work. Inspection of the output probability maps occasionally reveals misclassifications, particularly at the borders of the tissue microarray spots. Most misclassifications at the tissue borders are due to tissue preparation artefacts, which are not recognized by the network. To remedy this situation in a clinical application, an additional neural network could be trained to detect artefact regions and exclude them as a preprocessing step.

A more serious limitation, that is however not unique to our study, is the subjective nature of the Gleason scoring system. Inter-pathologist variability regarding Gleason score annotations is non-negligible, as also shown in the current study. Consensus annotations from multiple pathology experts would enable more objective training of an automated Gleason annotation model. Furthermore, disagreement in Gleason score annotations is expected to be even higher among pathologists from different hospitals, adhering to slightly different annotation guidelines and habits. Larger scale studies involving multiple medical centers are thus necessary to further consolidate and potentially improve our patient stratification results, and build a system that could be employed in clinical practice.

Furthermore, in current clinical practice, patients are stratified according to a novel prostate cancer grading system^37^ which is based on Gleason grading into five Grade Groups (corresponding to Gleason ≤ 6, Gleason 3+4, Gleason 4+3, Gleason 8, Gleason 9-10), each with distinct prognosis characteristics. However, this more fine-grained stratification is typically based on histological assessment of tissue areas larger than the TMA spots used in this study. In future work, we will evaluate our approach on whole slide tissue images with accompanying survival information.

Despite the limitations described above, the results of this study demonstrate the potential of the use of deep learning technology as assistance to pathologists in grading prostate cancer. Our study was performed on H&E stained tissue microarray spots, achieving pathology expert-level performance by learning from a remarkably small training cohort of 641 TMA spots. However, the approach is not specific to H&E staining and, therefore, similar strategies can be applied to more specific stainings for studying the effect of gene expression, tumor microenvironment or drug uptake on clinical outcome. Furthermore, in the presence of a larger cohort annotated with survival information, a deep neural network could be trained directly on survival endpoints, enabling the potential discovery of novel survival-associated morphological patterns and, ultimately, guiding towards the definition of a more objective prostate cancer grading scheme.

## Methods

### Tissue Microarray Gleason annotation

Tissue microarrays were digitized at 40x resolution (0.23 microns per pixel) at the University Hospital Zurich (NanoZoomer-XR Digital slide scanner, Hamamatsu). Tumour stage and Gleason scores were assigned according to the International Union Against Cancer (UICC) and WHO/ISUP criteria. Cancerous regions were delineated and labeled with corresponding Gleason grades using the TMARKER^38^ software. In addition, TMA spots containing only benign prostate tissue were marked as “benign” by the two pathologists. Use of image and clinico-pathological data of each cohort was approved by the Zurich Cantonal Ethics Committee (KEH-ZH-No. 2014-0604, 2007-0025, 2008-0040, 2014-0007). Clinico-pathological data have already been published in part by Zhong *et al*.^27^.

### Automated tissue detection

Tissue regions were automatically detected in each TMA spot by the following pipeline. Gaussian filtering was initially performed to remove noise, followed by Otsu thresholding that separated tissue from background. The tissue mask was further refined via morphological operations (dilation to fill in little holes in the tissue, followed by erosion to restore borders). Detected tissue regions were subsequently used to extract tissue patches and compute pixel-level probability maps.

### Patch creation

The original image resolution of individual TMA spots was 3100 x 3100 pixels. For model training, small image regions of size 750 x 750 were sampled from each TMA spot, using a step of 375 pixels. We also experimented with smaller image patches of size 300 x 300, but achieved inferior results on the validation set. Each patch was labeled according to the annotation in its central 250 x 250 region. Patches containing no or multiple annotations in the central region were discarded.

### Training details and model selection

We first conducted a model selection experiment, during which we evaluated training neural networks from scratch or fine-tuning them starting from ImageNet-learned parameters. The model variants considered were VGG-16, Inception V3, ResNet-50, MobileNet and DenseNet-121.

We only used the convolutional part of each model’s architecture, removing all fully-connected layers. On top of the last convolutional layer, we added a global average pooling layer, followed by the final classification layer that uses softmax non-linearity. For training from scratch, we used the Adam^39^ optimization technique with initial learning rate of 0.001. For fine-tuning the networks, we used SGD with learning rate 0.0001 and Nesterov momentum 0.9. DenseNet and MobileNet were trained with dropout of 0.2. In all cases, the categorical cross-entropy loss was used as minimization objective function.

Image patches were initially resized to 250 x 250. Data augmentation was applied during training to combat overfitting. We performed random cropping of 224 x 224 regions followed by random rotations, flipping and color jittering. Training with balanced mini-batches was crucial for achieving good validation performance across all classes. We used mini-batches of size 32 and, at each iteration, an equal number of examples from each class was randomly selected. All networks were trained for 50’000 iterations.

We used three tissue microarrays for model training (TMAs 111, 199, 204) and one for validation (TMA 76). The results of this experiment are depicted in **Supplementary Figure S1**. We observed that the networks trained from scratch converged in the first 20’000 iterations. Fine-tuned networks converged earlier and remained at lower validation loss levels than their randomly-initialized counterparts. We also noticed that several model variants converged to a validation accuracy close to 65% but at the same time exhibited increasing validation loss curves. This contrasting behaviour was more pronounced in the networks trained from scratch and can be explained by the following argument: networks trained from scratch (or lacking appropriate regularization) perfectly overfit to the training set and, thus, become very confident in all their predictions, producing higher cross-entropy loss in case of misclassifications. On the contrary, fine-tuned networks that are additionally regularized by dropout, such as MobileNet and DenseNet, do not reach 100% accuracy in the training set and do not exhibit an explosion in the validation cross-entropy loss scores. The best validation loss trajectory was achieved by the MobileNet network (using width multiplier *α* = 0.5) and, therefore, MobileNet was selected for further evaluation on the test cohort.

### Selected model architecture

The MobileNet^33^ network architecture is built as a series of “depthwise separable” convolution blocks. A depthwise separable convolution block consists of 3×3 convolutions applied *separately* to each input channel (depthwise), followed by 1×1 (pointwise) convolutions that combine the depthwise convolution outputs to create new features. Depthwise convolutions are occasionally performed with a stride *s* = *2* to reduce the dimensionality of the output feature maps. The coupled pointwise convolutions are then always doubling the number of output channels. A full MobileNet network starts with 32 channels at the first convolutional layer and increases up to 1024. The width multiplier parameter *α* can be used to build thinner networks, having even fewer parameters. In this study we used *α* = 0.5, thus starting with 16 channels and increasing up to 512. After the convolution blocks, a global average pooling layer is used to compute the spatial average of each feature map at the last convolution layer. Finally, the output layer uses a softmax activation function to compute an output probability distribution over the four Gleason classes considered in this study.

### Evaluation on test cohort

#### Choice of metrics

Throughout the manuscript, we use two metrics for quantifying inter-annotator variability. The first one is Cohen’s kappa statistic^34^, which is widely used for measuring inter-rater agreement. Cohen’s kappa takes into account the possibility of agreement occurring by chance, resulting in a value of 0 for agreement occurring by chance and a value of 1 for perfect agreement. For ordered classes, weighted Cohen’s kappa is more appropriate because it penalizes more strongly the inter-annotator disagreement occurring between more distant classes. Here, we use the quadratic weighted kappa statistic defined as follows:

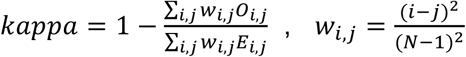

where *N* is the total number of considered classes or rating scores and the indices *i, j* refer to the ordered rating scores 1 ≤ *i, j* ≤ *N*. *O*_*i*_,_*j*_ denotes the number of images that received rating score *i* by the first expert and rating score *j* by the second and *E*_*i*_,_*j*_ denotes the expected number of images receiving rating *i* by the first expert and rating *j* by the second, assuming no correlation between rating scores.

In addition, we use macro-average recall as a metric that receives equal contribution from all classes, irrespective of class imbalance. Macro-average recall is computed as the unweighted average of recall values for individual classes.

#### Patch-level evaluation

For the confusion matrices in **Figure 2**, we only included patches annotated by both pathologists, so that model-versus-pathologist and inter-pathologist variability are quantified on the same set of examples.

#### TMA spot-level evaluation

To compute pixel-level output probability maps (**Fig. 3**), the network was converted to an equivalent fully convolutional architecture. The global average pooling layer, which acted on a 7 x 7 dimensional input when training on patches, was replaced with a local 7-by-7 average pooling layer with stride *s* = 1. The final classification layer was replaced with an equivalent convolutional layer with four output channels, one for each Gleason class. Finally, an upsampling layer with factor 32 was used to restore the dimensions of the input image and produce the pixel-level output probability maps.

The network-based composite Gleason score was assigned on the basis of the predicted probability maps. Let 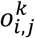 be the predicted probability for class k in location (i, j). An initial weighted score was computed for each class as 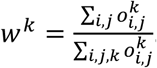. The final score was assigned to each class as 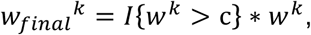 where *I* denotes the indicator function and *c* is a predefined threshold value. We set *c* = 0.25 to ensure that at least one class has a non-zero final score, since there are four output classes considered. The primary and secondary Gleason patterns were assigned on the basis of the final scores for each class and the composite Gleason score was computed as the sum of the primary and secondary patterns.

#### Model interpretation

Patches predicted with high confidence (probability > 0.8 for the correct class) were pre-selected and visualized. Within these high-confidence patches, we seeked to identify class-specific discriminative subregions. To this end, we applied the class activation mapping (CAM) technique^35^, which is particularly suitable for networks with a fully convolutional architecture and a global average pooling layer immediately before the final classification layer. Class activation maps are generated by projecting the class-specific weights of the output classification layer back to the feature maps of the last convolutional layer, thus highlighting important regions for predicting a particular class.

## Code availability

Model training and evaluation were performed in Python 3 using the deep learning library Keras with Tensorflow backend. Survival analysis was performed in R using the packages `survival` and `survminer`.

The scripts are available on Github (https://github.com/eiriniar/gleason_CNN).

## Data availability

All tissue microarray images used in this study will be made publicly available upon publication, together with corresponding Gleason annotations provided by the pathologists. Patient survival data have been previously published by Zhong *et al*.^27^ and are available upon request from the authors of that study^27^.

## Acknowledgements

This work was sponsored in part by a grant from the Swiss National Science Foundation (zBioLink), a SystemsX.ch grant (Phosphonet-Personalized Precision Medicine) and a grant provided by the Foundation for Research in Science and the Humanities at the University of Zurich provided to PJW.

## Author contributions

EA and MC designed the computational experiments. EA performed the computational experiments, analyzed the data and wrote the manuscript. KSF provided Gleason annotations for all TMAs and wrote the manuscript. MM contributed to computational experiments. PJW, NJR, TH, CF and NW contributed to TMA and clinical data aquisition. JHR supervised the study, provided Gleason annotations for the test cohort and wrote the manuscript. MC and PJW conceived, designed and supervised the study, and wrote the manuscript. All authors edited and approved the manuscript.

